# Extracellular DJ-1 induces sterile inflammation in the ischemic brain

**DOI:** 10.1101/2020.09.16.299420

**Authors:** Koutarou Nakamura, Seiichiro Sakai, Jun Tsuyama, Akari Nakamura, Kento Otani, Kumiko Kurabayashi, Yoshiko Yogiashi, Hisao Masai, Takashi Shichita

## Abstract

Inflammation is implicated in the onset and progression of various diseases, including cerebral pathologies. Here we report that DJ-1, which plays a role within cells as an antioxidant protein, functions as a damage-associated molecular pattern (DAMP), and triggers inflammation if released from dead cells into the extracellular space. We first found that recombinant DJ-1 protein induces the production of various inflammatory cytokines in bone marrow-derived macrophages (BMMs). We further identified a unique peptide sequence in the αG and αH helices of DJ-1 that activates Toll-like receptor 2 (TLR2) and TLR4. In the ischemic brain, DJ-1 is released into the extracellular space from necrotic neurons within 24 hours after stroke onset and makes direct contact with the surfaces of infiltrating myeloid cells. Administration of an antibody against DJ-1 suppresses the expression of inflammatory cytokines in infiltrating immune cells and attenuates ischemic neuronal damage. Our results demonstrate a previously unknown function of DJ-1 as a DAMP and suggest that extracellular DJ-1 could be a therapeutic target to prevent inflammation in tissue injuries and neurodegenerative diseases.

**Significance statement:** DJ-1 has been thoroughly investigated as a cytoprotective antioxidant protein in neurons. However, here we demonstrate that extracellularly released DJ-1 triggers neurotoxic inflammation after ischemic stroke. Intracellular DJ-1 increases in response to oxidative stress in ischemic neurons, but if ischemic stresses result in necrotic cell death, DJ-1 is released extracellularly. Released DJ-1 interacts with TLR2 and TLR4 on the surface of infiltrating myeloid cells and triggers post-ischemic inflammation, leading to the exacerbated pathologies of ischemic stroke. Thus, extracellular DJ-1 is a previously unknown inflammatogenic DAMP, and may be a putative target for therapeutic intervention to prevent progression of inflammatory and neurodegenerative diseases.

## Introduction

Inflammation is implicated in the pathophysiology of a wide range of illnesses, including neurological diseases and psychiatric disorders (Pape *et al*, 2019). Circulating or tissue-resident immune cells trigger inflammation upon encountering stimuli such as infections and tissue damage (Fan & Rudensky, 2016). During infections, pathogen-specific compounds induce the production of various inflammatory mediators in innate immune cells through the activation of pattern recognition receptors (PRRs) (Kawai & Akira, 2010). PRRs are also activated by endogenous ‘self’ molecules released into the extracellular space from damaged tissue (Kono & Rock, 2008). These endogenous alarm molecules trigger sterile inflammation and are known as DAMPs. Since the brain is generally a sterile organ, cerebral inflammation is thought to be induced by DAMPs released from damaged brain cells (Chen & Nuñez, 2010). Recently, the prevention of excess inflammation has been reported to improve the symptoms of neurological diseases and psychiatric disorders (Iadecola & Anrather, 2011). Controlling cerebral inflammation by targeting DAMPs could be a promising therapeutic method for cerebral pathologies.

Ischemic stroke, which is a major cause of death and disability all over the world, is the sudden onset of neurological deficits due to necrosis of brain tissue caused by a severe loss of cerebral blood flow (CBF). The presence of large quantities of DAMPs generated by necrotic brain tissue triggers severe inflammation, which worsens the neurological deficits associated with stroke (Dayon *et al*, 2011). DAMPs are thus key molecules in determining an individual’s functional prognosis after ischemic stroke. Among the various PRRs that recognize DAMPs, TLR2 and TLR4 are pivotal in cerebral post-ischemic inflammation, given that TLR2- and TLR4-deficiencies considerably decrease the expression of various inflammatory mediators in the ischemic brain (Shichita *et al*, 2009). In ischemic stroke, high mobility group box 1 (HMGB1) and the peroxiredoxins (PRXs) have been identified as the DAMPs (Qiu *et al*, 2008; Shichita *et al*, 2009). HMGB1 is a DAMP with an immediate effect, as it is extracellularly released within six hours after stroke onset and promotes disruption of the blood-brain barrier (Qiu *et al*, 2008). The PRXs, on the other hand, trigger the expression of various inflammatory cytokines including IL-23 in infiltrating macrophages through the activation of TLR2 and TLR4 (Shichita *et al*, 2012). IL-23 from infiltrating macrophages induces the expression of IL-17 in γδT lymphocytes, which promotes post-ischemic inflammation in the delayed phase of ischemic stroke (Gelderblom *et al*, 2012; Shichita *et al*, 2009). PRXs identified from brain lysates have been shown to induce the expression of inflammatory cytokines in cultured myeloid cells (Shichita *et al*, 2012). This induction is partially decreased by the depletion of PRX proteins in brain lysates.

In this study, we searched for previously unknown DAMPs in brain lysates. We identified DJ-1 (alternatively known as PARK7) as a major DAMP with a unique peptide sequence that activates TLRs.

## Results

### DJ-1 induces the production of inflammatory cytokines in BMMs through TLR2 and TLR4 activation

We previously found prominent DAMP activity in the 15 to 25 kDa fractions of brain lysates (Shichita *et al*, 2012). Among candidate DAMPs detected in these fractions by mass spectrometry, we generated the recombinant proteins, and added these individually to a culture of BMMs to examine DAMP activity. We discovered that recombinant DJ-1 protein induced mRNA expression of inflammatory cytokines such as TNFα, IL-1β and IL-23 (a heterodimer of IL-23p19 and IL-12p40) in a dose-dependent manner (**Fig.1A**), while a control protein, GST (glutathione-S-transferase), did not. mRNA expression of TNFα, IL-1β and IL-23 increased within 6 hours after the addition of DJ-1 protein in BMMs (**Fig.1B**). Amounts of TNFα and IL-23 protein in the culture supernatant of BMMs increased three hours after the stimulation by DJ-1, which were significantly higher than GST-treated BMMs (**Fig.1C**).

**Figure 1.**
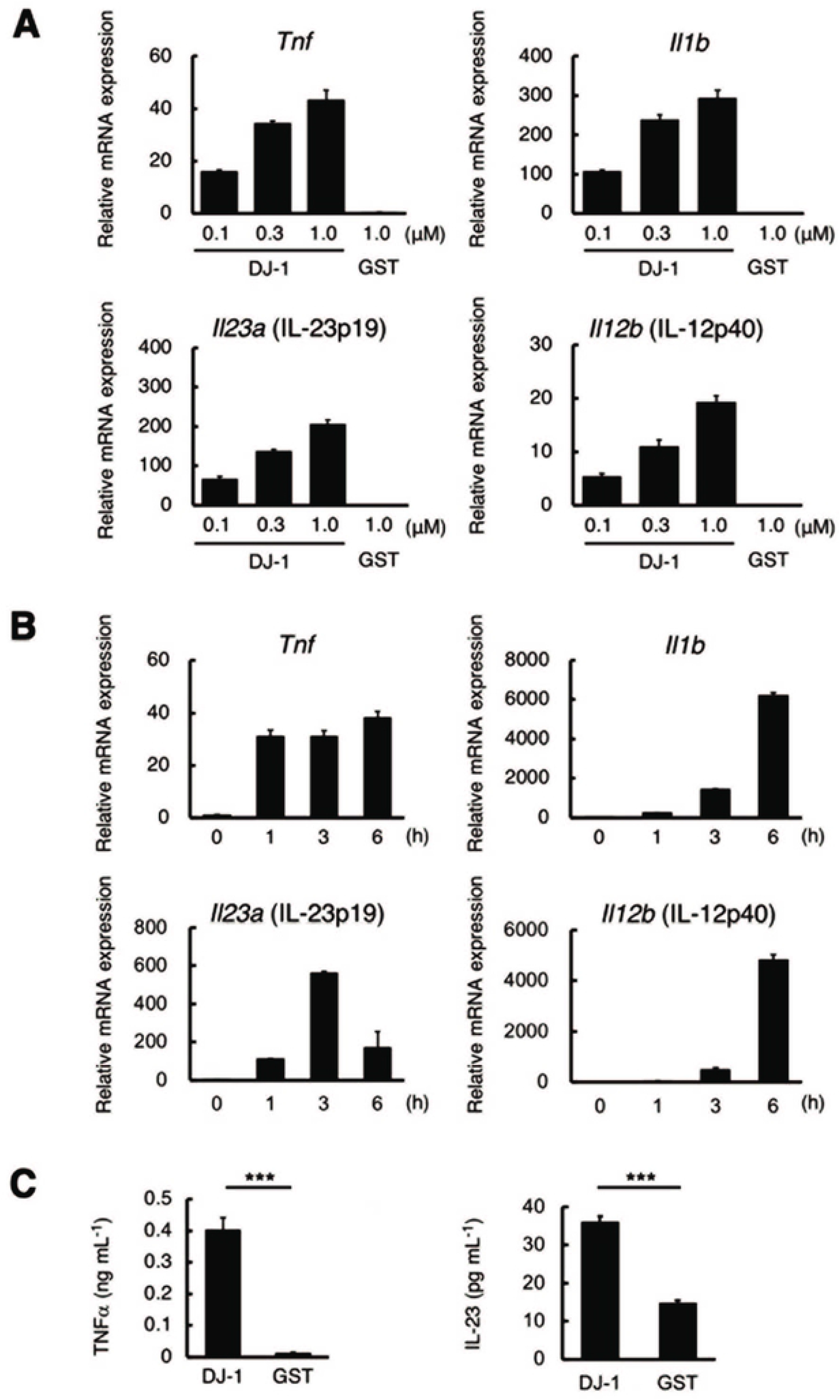
DJ-1 induces the expression of inflammatory cytokines in BMMs. **(A)** The mRNA expression levels of inflammatory cytokines in BMMs treated with the indicated concentrations of recombinant DJ-1 or GST protein for 1 h (*n* = 3 for each group). Each value indicates the relative mRNA expression evel compared to that of *Hprt1.* **(B)** Time-dependent changes in mRNAexpression level of inflammatory cytokines in BMMs treated w th 1 μM of recombinant DJ-1 protein (*n* = 3 for each group). **(C)** TNFα and IL-23 protein expression levels in the BMM supernatant 3 hour s after treatment with 1 μM of recombinant DJ-1 or GST protein (*n* = 3 for each group). \*\*\**P* < 0.001 vs. GST-treated BM Ms **(C)** (two-sided Student’s *t*-test **[C]**). The error bars represent s.e.m.

Since TLRs are major PRRs which mediate the inflammatory responses of various DAMPs, we next explored whether this is also the case for DJ-1 by HEK293 cells expressing each TLR family member. DJ-1-induced activation of nuclear factor kappa B (NFκB) was investigated using a luciferase reporter assay in these HEK293 cells. Recombinant DJ-1 protein induced luciferase activity in TLR2-expressing HEK293 cells but not in control HEK293 cells (**Fig.2A**). A positive control, peptidoglycan (PGN), also induced luciferase activity in TLR2-expressing HEK293 cells. A similarly increase in luciferase activity was also observed by the addition of recombinant DJ-1 protein to TLR4/MD-2/CD14-expressing HEK293 cells, comparable to the increase induced by stimulation with lipopolysaccharide (LPS) (**Fig.2B**). Control GST protein did not induce luciferase activity in any of these cell types. Although we also examined the activation of endosomal TLRs (TLR3, TLR7, or TLR9) by recombinant DJ-1 or intracellular overexpression of DJ-1, we could not detect any luciferase activity in these endosomal-TLRs-expressing HEK293 cells (**Fig.2C** **and Fig.S1**). Therefore, we identified TLR2 and TLR4 as PRRs for DJ-1, and this was confirmed by using BMMs generated from TLR2 and TLR4 double-deficient mice. As shown in **Fig.2D**, the expression of inflammatory cytokines induced by DJ-1 was almost completely abolished in TLR2 and TLR4 double-deficient BMMs. Thus, both TLR2 and TLR4 are PRRs for the DJ-1 protein to trigger inflammatory responses.

**Figure 2.**
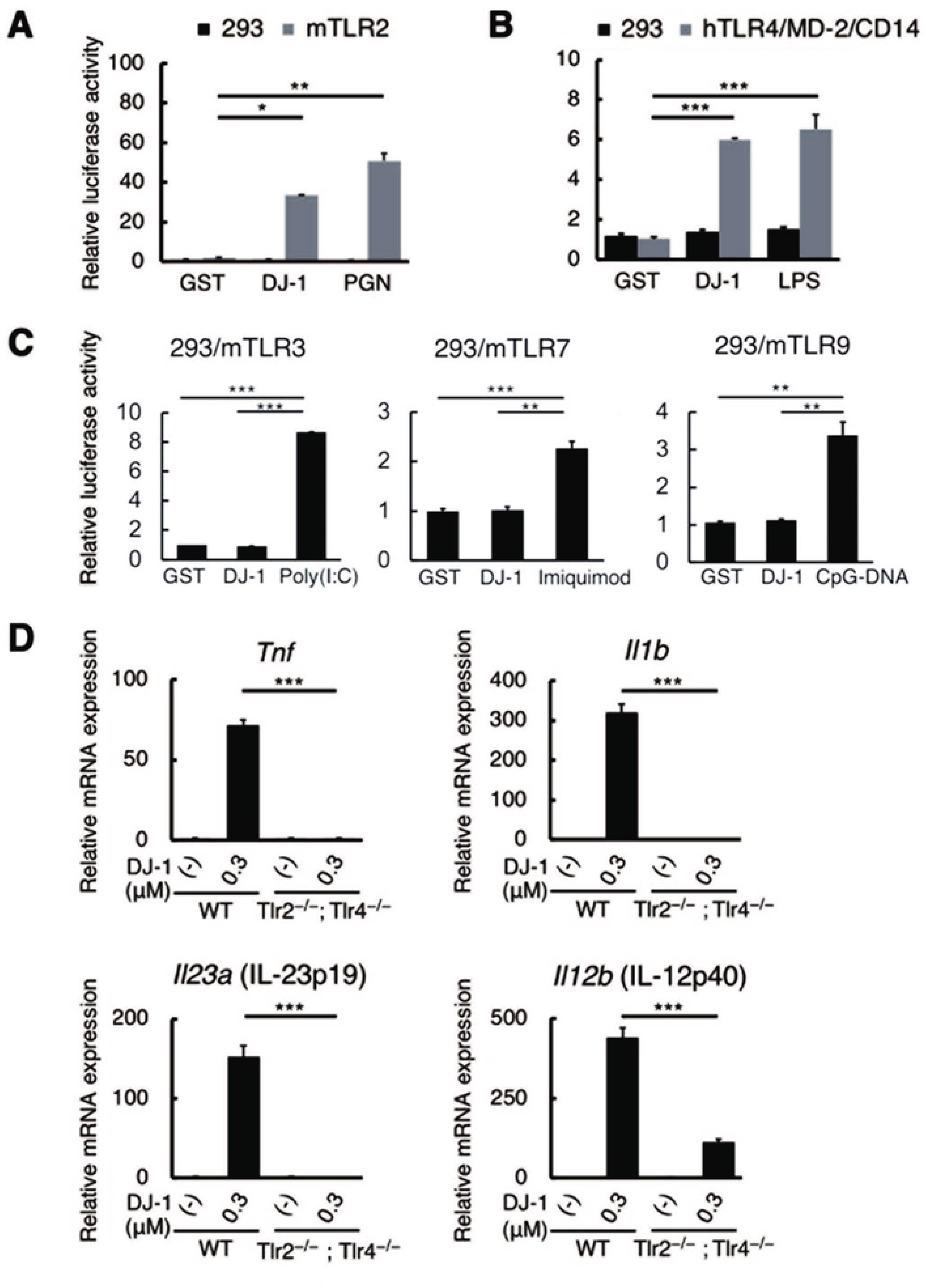
DJ-1 activates TLR2 and TLR4 to trigger inflammation. **(A)** The relative luciferase activity of NF-κB reporter in HEK293 cells or mur ne stably TLR2-expressing HEK293 cells treated with 1 μM of recombinant DJ-1 protein or PGN (TLR2 ligand) *(n* =3 for each group). **(B)** The relative luciferase activity of NF-κB reporter in HEK293 cells or human stab y TLR4/MD-2/CD14-expressing HEK293 cells treated with 1 μM of recombinant DJ-1 protein or LPS (TLR4 ligand) (*n* = 3 for each group). The relative val ues compared to those in HEK293 cells treated with 1 μM of contro GST prote in are shown **(A,B)**. **(C)** Relat ve luciferase activity of the NF-KB reporter in each mur ne TLRs expressing HEK293 cells treated w th 1 pM of recombinant GST (as a control) or DJ-1 protein or TLR ligands (Poly(I:CJ): a TLR3 ligand. lmuquimod: a TLR7 ligand. and CpG-DNA: a TLR9 ligand). **(D)** The mRNA expression levels of inflam matory cytokines in WT or TLR2 and TLR4 doub e deficient BMMs 1 h after stimulation with recombinant DJ-1 protein (*n* = 3 for each group). The relative values compared to those in untreated WT BMMs are shown. \**P*<*0.05*. \*\**P*<*0.01*, \*\*\**P* < 0.001 (one-way ANOVA with Dunnetrs correction **[A,B,C,D]**). The error bars represent s.e.m.

### The αG and αH helix region of DJ-1 is important for DAMP activity

Since DJ-1 is known to be an antioxidant protein whose cysteine 106 (Cys-106) is important for reducing reactive oxygen species (ROS) (Andres-Mateos *et al*, 2007), we generated a mutant DJ-1 protein which lacks the ability to form the oxidized state, or dithiothreitol-treated DJ-1 protein. However, these modifications did not affect DAMP activity (**Fig.S2**), suggesting that DAMP activity and antioxidant activity are separable functions of DJ-1. Next, in order to identify the peptide sequence in DJ-1 required for activation of TLR2 and TLR4, we generated various DJ-1 deletion mutant peptides fused with GST (**Fig.3A**). We examined IL-23-inducing activities of C-terminal deletion mutants using BMMs and found that a deletion between residues 160 and 189 of DJ-1 significantly decreased IL-23-inducing activity (**Fig.3B**). Thus, we focused on the C-terminus region of DJ-1. GST-fused to residues 100 to 189 of DJ-1 induced IL-23 to a level similar to full-length DJ-1 protein (**Fig.3C**). While residues 100 to 160 did not induce IL-23 expression in BMMs, induction by residues 160 to 189 was again similar to that of full-length DJ-1 protein (**Fig.3C**). This region is located at the surface of the DJ-1 molecule and contains the αG helix and αH helix regions (**Fig.3D**). Although DJ-1 is an atypical peroxiredoxin-like peroxidase, the peptide sequence between residues 100 and 120 of DJ-1, which exhibits no DAMP activity, is most similar to the peptide sequence in the β4 sheet and α3 helix regions of PRXs, which are important for the DAMP activities exhibited by the PRXs (Shichita *et al*, 2012) (**Fig.3E**). As we found no similarity between the peptide sequence of the αG–αH helix region of DJ-1 and that of any other proteins bearing DAMP activity (HMGB1, PRXs, S100A8/A9, etc.), DJ-1 seems to have a unique peptide sequence that triggers the production of inflammatory cytokines in BMMs.

**Figure 3.**
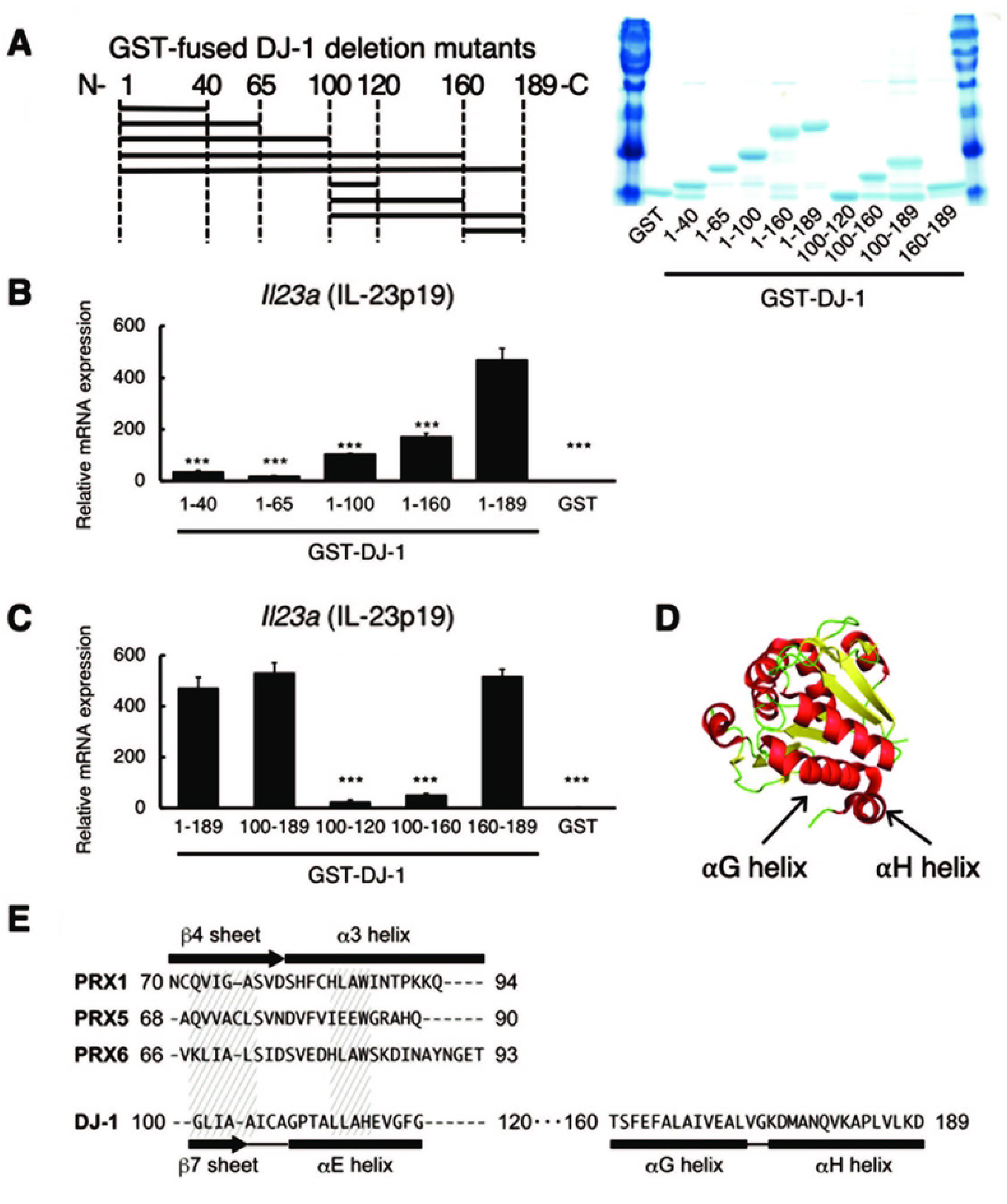
The αG and αH helix region of OJ-1 is essent al for DAMP activity. (**A**) Diagram of generated deletion mutant peptides of DJ-1 fused with GST (left panel). The number of amino acid residues contained in each GS Tfused DJ-1 peptide is shown on the x axis. The amount of generated deletion mutant peptides was examined by SOS PAGE with CBB staining (right panel). (**B**) IL 23p19 inducing ac tivities of C terminal deletion mutants of OJ-1. (**C**) IL 23p19 inducing activ ties of GSTfused peptides containing the C-terminal region of DJ-1. (**D**) αG helix (DJ-1_160-173_: peptides between residues 160 and 173 of DJ-1) and αH helix (DJ-1_175-189_) are indicated in the crystal structure of DJ-1 protein (PDB ID: 1PSF). (**E**) Compar son of peptide sequences among PRX1_70-94_. PRX5_68-90_. and P RX6_66-93_ containing the β4 sheet and α3 helix of PRXs known to be important for DAMP activity. DJ-1_100-120_ containing the β7 sheet and aE helix. and DJ-1_160-189_ containing the αG and αH helix. Similar ties among the amino acid res dues of PRXs and DJ-1 are indicated by hatched areas. \*\*\**P*< 0.001 vs. BMMs treated with DJ-1_1-189_ (**B,C**) (one-way ANOVA with Dunnett’s correction [**B,C**]). The error bars represent s.e.m. and *n* = 3 for each group (**B,C**).

### DJ-1 is extracellularly released from necrotic brain cells and stimulates infiltrating macrophages

Next, we examined the function of extracellular DJ-1 as a DAMP in the ischemic brain using a murine model of brain ischemia-reperfusion. In the normal brain, DJ-1 protein could be detected in neurons, astrocytes, oligodendrocytes, and myeloid cells using tyramide-mediated signal amplification (**Fig.S3A**). DJ-1 was not detected in normal, non-ischemic neurons using non-enhanced staining in the absence of tyramide, but was detected in ischemic neurons 6 to 12 hours after stroke onset (**Fig.4A**), revealing that ischemic stress induces DJ-1 expression in neurons. Twenty-four hours after stroke onset, NeuN-positive cells did not survive in the infarct area and DJ-1 was detected in debris-like structures. Staining with control IgG did not yield any signals in ischemic brains (**Fig.4A**). The specificity of our anti-DJ-1 antibody was confirmed by the detection of a single DJ-1 band in lysates from ischemic brain tissue by Western blotting (**Fig.S3B**). Around the infarct boundary zone, DJ-1 expression was observed within NeuN-positive cells even 24 hours after stroke onset (**Fig.4B**); in contrast, in the infarct area, cell debris staining positive for DJ-1 was observed separate from cellular membranes (stained with pan-cadherin antibody) (**Fig.4C**). This extracellular DJ-1-including debris was also observed even when the ischemic brain tissue was not re-perfused, indicating that the reperfusion after brain ischemia is not necessary for the extracellular release of DJ-1 (**Fig.S4A,B**). Debris containing DJ-1 was observed scattered around TUNEL-positive necrotic brain cells in the infarct region 24 hours after stroke onset (**Fig.4D**). We also found that DJ-1-including debris was in direct contact with the F4/80-positive cellular membranes of infiltrating myeloid cells in the infarct region (**Fig.4E,F**). These observations suggest that DJ-1 is released into the extracellular space from necrotic brain cells where it directly activates infiltrating myeloid cells.

**Figure 4.**
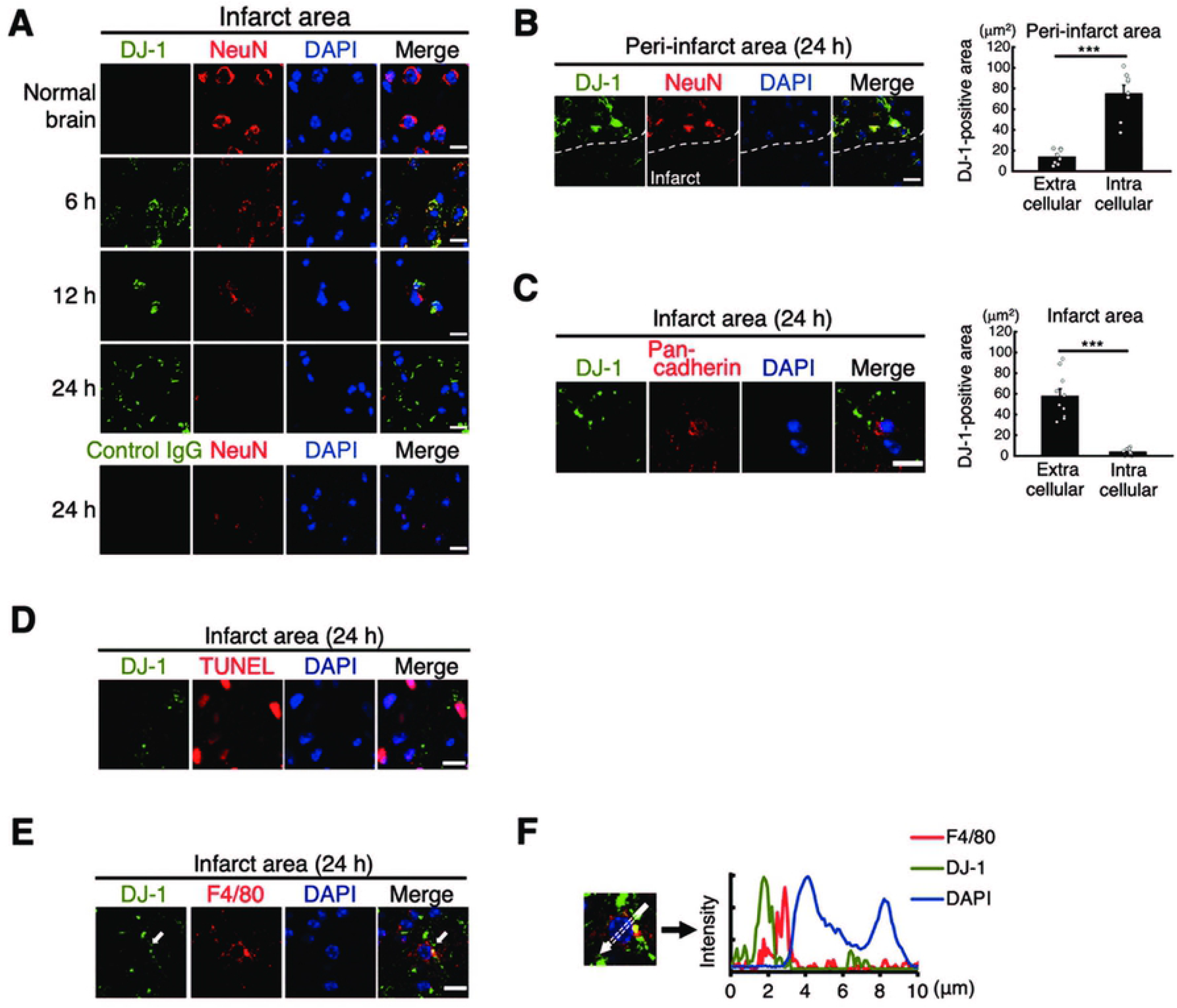
DJ-1 is released into the extracellular space from necrotic brain cells and contacts infiltrating myeloid cells in the ischemic brain. (**A**) lmmunohistochemical staining of DJ-1. NeuN and DAPI in normal and ischemic brain. Time-dependent changes in infa rct area were investigated at the indicated time points. Specificity of our staining technique for DJ-1 was confirmed by testing with control lgG. **(B)** lmmunohistochemical staining of DJ-1. NeuN and DAPI in the peri infarct area 24 h after stroke onset. Dashed white line indicates infarct border (left panel). Quantification of intracellular or extracellular DJ-1-posit ve areas in the peri-infarct region 24 h after the stroke onset (right panel. *n* = 8 for each group). (**C**) lmmunohistochemica l staining of DJ-1. pan cadherin and DAPI in the infarct region 24 h after the stroke onset (left panel). Quan tification of intracellular or extracellular DJ-1-positive areas in the infarct region 24 h after the stroke onset (right panel. *n* = 10 for each group) **(D)** lmmunohistochemical staining of DJ-1 in the infarct region 24 h after stroke onset. Dead brain cells were detected by TUNEL staining. (**E**) lmmunohistochemical staining of DAPI. F4/80 and DJ-1 in the infarct region 24 h after stroke onset. White arrow indicates direct contact of DJ-1-including debris with the cellular membranes of infiltrating myeloid cells. **(F)** Fluorescence intensity along the white arrow shown in **Fig.4E**. (scale bars: 10 μm (**A,B,C,D,E**)). All images were captured using confocal laser microscopy. (two-sided Student’s *t*-test [**B,C**]) \*\*\**P*<*0.001* vs. extracellular DJ-1-positive area (**B.C**). The error bars reoresent s.e.m.

### DJ-1 functions as a DAMP in ischemic stroke

To prove our hypothesis that the activation of infiltrating myeloid cells by DJ-1 leads to the progression of ischemic neuronal injury, we examined whether the deficiency of DJ-1 or the administration of a DJ-1-specific antibody exerted an immunosuppressive effect in the murine model of ischemic stroke. We found that DJ-1-deficiency significantly decreased the expression of inflammatory cytokines such as TNFα, IL-1β, and IL-23p19 in the infiltrating immune cells collected from ischemic brain (**Fig.5**). Consistent with the previous report demonstrating the neuroprotection by anti-oxidative effect of intracellular DJ-1 (Aleyasin et al 2007), the significant difference of infarct volume was not observed between WT and DJ-1-deficient mice when brain ischemia-reperfusion was induced by middle cerebral artery occlusion (data not shown).

**Figure 5.**
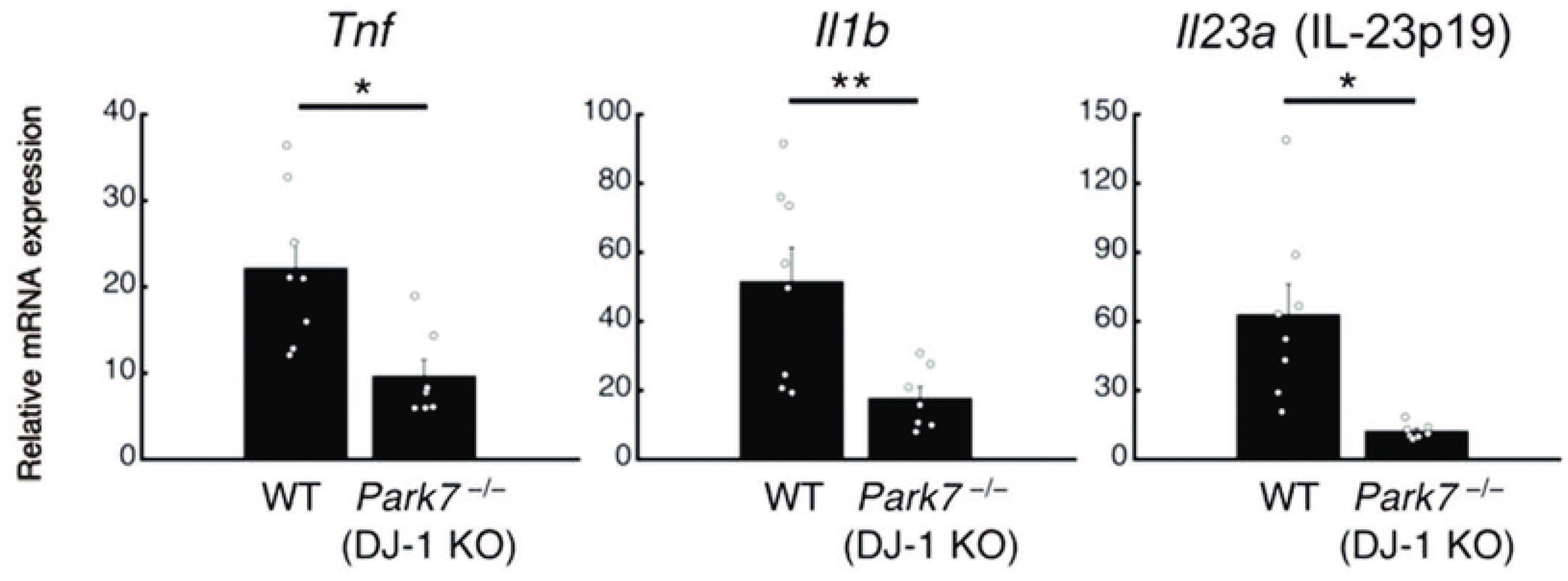
The expression of inflammatory cytokines was attenuated in DJ-1-deficient mice. The mRNA expression levels of inflammatory cytokines in infiltrating immune cells collected from day 1 post-ischemic brains of WT or DJ-1 KO mice (*n* = 8 for WT. *n* = 7 for DJ-1 KO). The relative values compared to those in sham-operated mice are shown. \**P*<*0.05*. \*\**P*<*0.01* vs. WT mice (two-sided Student’s t-test). Error bars represent the s.e.m.

We next examined the administration of a DJ-1-specific antibody to neutralize extracellular DJ-1. Compared to control IgG antibody, the DJ-1-specific antibody significantly decreased mRNA expression of IL-23p19 in infiltrating immune cells 24 hours after stroke onset (**Fig.6A**). mRNA expression of TNFα and IL-1β in ischemic brain tissue on day 3 after stroke onset was also decreased in mice treated with anti-DJ-1 antibody (**Fig.6B**). The number of infiltrating myeloid cells was not altered by the administration of antibodies (**Fig.S5**). Therefore, the neutralization of extracellular DJ-1 did not inhibit the infiltration of immune cells but suppressed the neurotoxic inflammation in the ischemic brain tissue. Indeed, extracellular DJ-1 did not directly induce neuronal cell death but triggered the production of neurotoxic inflammatory mediators through the activation of macrophages (**Fig.S6**). Moreover, the reduction of inflammatory cytokine expression in infiltrating immune cells by the administration of anti-DJ-1 antibody was not observed if macrophages were depleted by clodronate liposome administration (**Fig.S7**).

**Figure 6.**
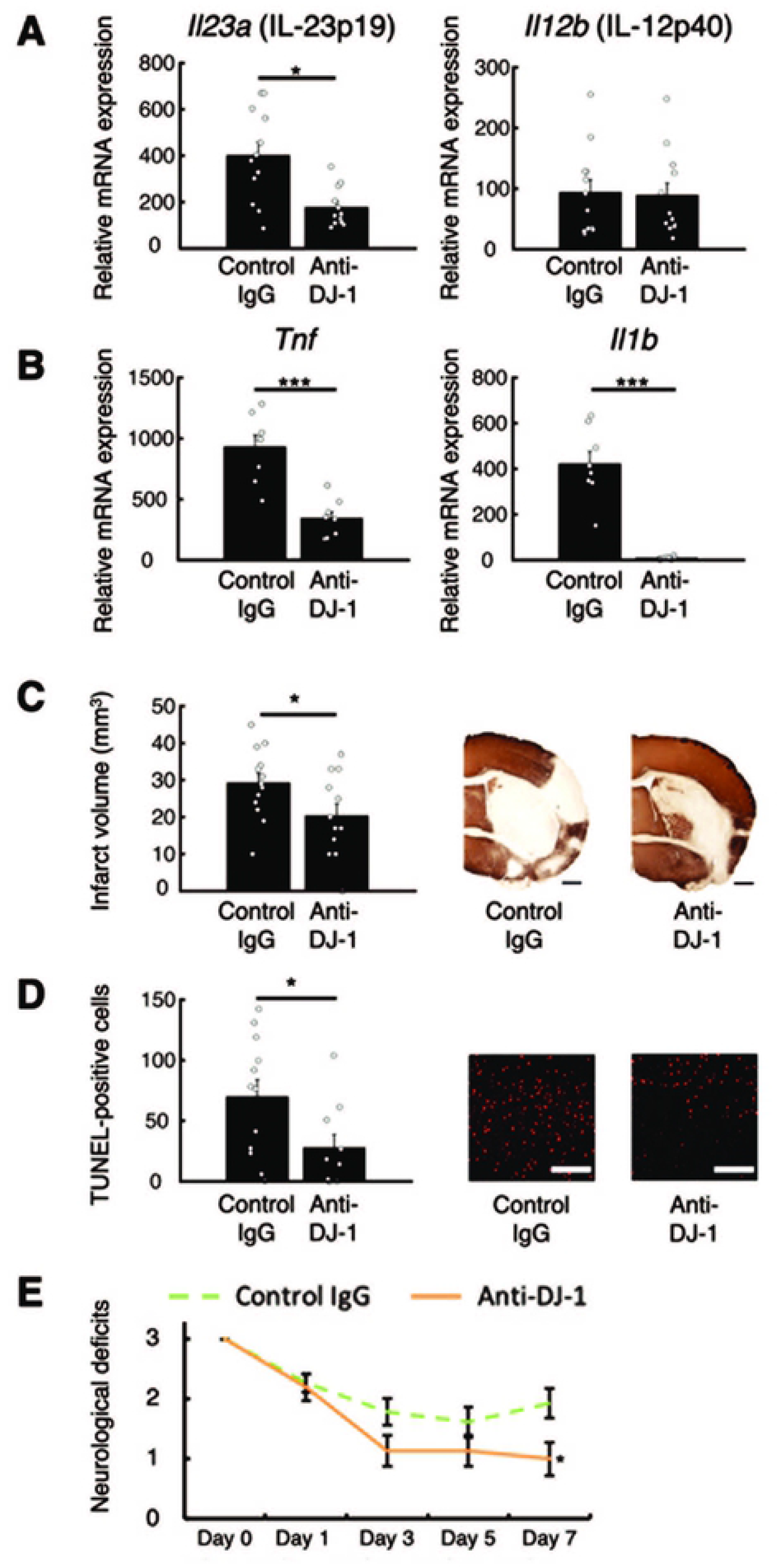
Administration of anti-DJ-1 antibody decreased the expression level of inflammato ry cytolines and attenuated ischemic neuronal injury. (**A**) The mRNA expression levels of IL-23 (p19 and p40) in infiltrat ng immune cells collected from day 1 post­ schem c brains of mice treated w th DJ-1-specific antibody or control lgG antibody immediately after stroke onset(*n* = 12 for each group). (**B**) The mRNA expression levels of TNFα and IL-β in the ischemic brain tissue on day 3 after stroke onset (*n* = 8 for each group).The relative values compared to those in sham oper ated mice are shown (**A,B**). (**C**) Infarct volume of mice treated with DJ-1specific antibody or control lgG antibody on day 7 after stroke onset (*n* = 13 for control lgG. *n* = 12 for anti-DJ-1 antibody). (*D*) The absolute number ofTUNEL-posi t ve neuronal cells in the peri-infarct area of mice treated with each ant body on day 7 after stroke onset (*n* = 12 for control lgG. *n* = 10 for anti DJ-1 antibody) (scale bars: 500 μm (C). 100 μm (**D**)). (**E**) Neurological deficits of mice treated with each antibody until day 7 after stroke onset. (*n* = 19 for control lgG. *n* = 15 for anti DJ-1 antibody). \**P*<*0.05*. \*\*\**P*<*0.001* vs. mice treated v1ith control lgG antibody. (**A,B,C,D,E**) (two-sided Student’s t-test [**A,B,C,D,**] Wilcoxon rank sum test [**E**]). The error bars represent s.e.m.

Finally, we examined the neuroprotective effect of anti-DJ-1 antibody against ischemic stroke. The administration of anti-DJ-1 antibody immediately after stroke onset significantly reduced the infarct volume compared to the administration of control IgG antibody (**Fig.6C**). There was no significant difference in CBF between the two groups, although a slight improvement in survival rate was seen in the mice treated with anti-DJ-1 antibody (**Table S1,S2**). The number of TUNEL-positive dead neuronal cells in the peri-infarct area on day 7 after stroke onset was significantly decreased by the administration of anti-DJ-1 antibody (**Fig.6D**). Consistent with this, the neurological deficits in the mice treated with anti-DJ-1 antibody were significantly improved on day 7 after stroke onset (**Fig.6E**). Altogether, our results indicate that DJ-1 is a DAMP molecule that promotes neuronal injury after ischemic stroke.

## Discussion

DJ-1 was first identified as an oncogenic protein and later recognized as a pivotal protein associated with the onset of familial Parkinson’s disease (Bonifati *et al*, 2003; Nagakubo *et al*, 1997). DJ-1-deficient mice are less effective at scavenging mitochondrial hydrogen peroxide in old age, indicating that DJ-1 is an atypical peroxiredoxin-like peroxidase that catalyzes ROS (Mullett *et al*, 2009). Increased expression levels of DJ-1 within brain cells are observed in chronic neurodegenerative diseases (Aleyasin *et al*, 2007). Thus, DJ-1 is believed to prevent neurodegeneration by reducing oxidative stress within brain cells. A neuroprotective role of intracellular DJ-1 against ischemic brain injury has been reported by using an endothelin-1 injection model (Aleyasin *et al*, 2007). In contrast, we did not observe significant differences in infarct volume between WT and DJ-1 KO mice (data not shown). This discrepancy is likely due to the different experimental methods used for the induction of brain ischemia. We used a middle cerebral artery occlusion (MCAO) model. A drastic infiltration of immune cells is generally observed in MCAO models. Thus, the neurotoxic effect of inflammation induced by extracellular DJ-1 should be greater, compared to using an endothelin-1 injection model. In our MCAO model, the neuroprotective effect of intracellular DJ-1 seems to be canceled by the inflammation-mediated neuronal damage triggered by extracellular DJ-1.

Our current study reveals that extracellularly released DJ-1 protein from necrotic brain cells triggers cerebral post-ischemic inflammation through the activation of TLR2 and TLR4 in infiltrating myeloid cells. We observed an increased expression of DJ-1 within neurons in the ischemic brain, suggesting that intracellularly accumulated DJ-1 is released into the extracellular space where it then functions as a DAMP. Thus, DJ-1 seems to have two opposing functions in the intracellular and extracellular contexts (**Fig.7**). Since important roles of DJ-1 have been demonstrated in several neurodegenerative diseases (Annesi *et al*, 2005; Bonifati *et al*, 2003; Glat *et al*, 2016; Lei *et al*, 2018; Lev *et al*, 2015), the function of extracellular DJ-1 as an inflammatory DAMP may be implicated in the pathophysiology of neurodegenerative diseases including Parkinson’s disease.

**Figure 7.**
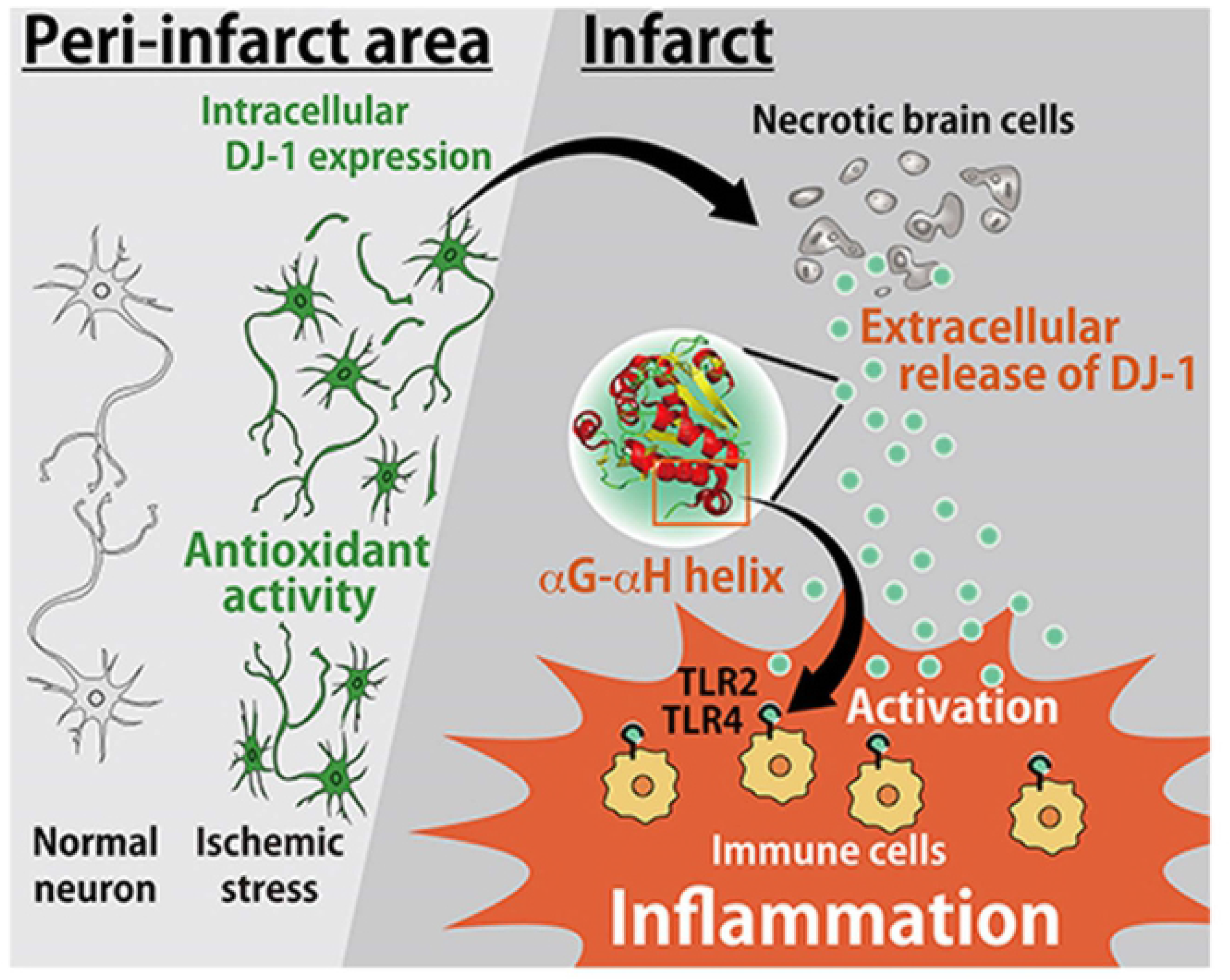
Schematic model of the roles of DJ-1 in the ischemic brain injury. Around the peri-infarct area. the expression level of DJ-1 within the neurons increases due to ischemic stress. Intracellular DJ-1 plays a neuroprotective role by catalyzing ROS. whereas if ischemic stress results in neuronal cell death, accumulated DJ-1 within ischemic neurons is released into the extracellular space. The αG-αH helix of extracellular DJ-1 directly activates TLR2 and TLR4 in infiltrating immune cells to trigger sterile inflammation. Therefore. DJ-1 has two opposing functions extracellularly (inflammatory molecule as a DAMP) and intracellularly (antioxidant activity) in the pathology of ischemic stroke.

DJ-1 protein is an atypical peroxiredoxin-like peroxidase; therefore, we compared the crystal structures and amino acid sequences between DJ-1 and PRX family proteins. Although PRXs have one or two cysteine residues which are conserved among PRX family proteins and are important for their anti-oxidant activities, DJ-1 also has two cysteine residues (Cys-53 and Cys-106) in humans and three cysteine residues (Cys-53, Cys-106 and Cys-121) in mice. Among these, the oxidation-sensitive cysteine-106 (Cys-106) of DJ-1 has been demonstrated to be essential for reducing oxidative stress within neuronal cells after ischemic stroke (Aleyasin *et al*, 2007; Andres-Mateos *et al*, 2007). Hydrogen bonding between glutamate-18 (Glu-18) and Cys-106 of DJ-1 catalyzes ROS by converting between their reduced and oxidized forms (Gupta & Carroll, 2014). Once DJ-1 is released into the extracellular space, however, its antioxidant activity is abolished as its cysteine residues are rapidly oxidized (Toohey, 1975). Crystal structure analysis reveals almost no differences between reduced and oxidized DJ-1, suggesting that the antioxidative function of DJ-1 does not significantly affect its DAMP activity (Gupta & Carroll, 2014). We identified αG–αH helix region of DJ-1 as an active center of DAMP activity which was the unique structure compared to PRX family proteins. The neutralization of PRX1, PRX2, PRX5, and PRX6 is necessary for the suppression of inflammatory cytokine production in ischemic stroke (Shichita *et al*, 2012), while the significant attenuation of inflammation was observed even by the neutralization of only DJ-1. This may be advantageous for developing the therapeutic method of ischemic stroke.

TLR2 and TLR4 are pivotal PRRs in cerebral sterile inflammation. DJ-1 can activate both TLR2 and TLR4, although the adaptor proteins MD2 and CD14 are necessary for TLR4 activation. The peptide sequence of DJ-1, especially in the αG–αH helix of DJ-1, is highly conserved between bacteria and mammals, suggesting that this conserved peptide sequence is commonly recognized by TLRs. A previous report has demonstrated that the αG–αH helix is necessary for the dimerization of the DJ-1 protein, which may enable its antioxidant activity (Tao & Tong, 2003). However, dimerization of DJ-1 protein seems unnecessary for the activation of TLR2 and TLR4, since even the GST-fusion αG–αH helix peptide induces almost the same level of DAMP activity as the full-length DJ-1 does. Furthermore, the timing of extracellular release of DAMPs is also important for the activation of infiltrating immune cells in the damaged tissue. The infiltration of macrophages and neutrophils becomes remarkable within 24 hours after stroke onset, and the extracellular release of DJ-1 in the ischemic brain coincides with this infiltration of immune cells. Among the inflammatory cytokines, the expression of IL-23 in the ischemic brain is especially dependent on the TLR2 and TLR4 signaling pathway. IL-23 not only promotes ischemic neuronal injury on day 1 but also sustains the inflammatory response several days after the onset of ischemic stroke (Shichita *et al*, 2009). This knowledge is consistent with our current finding that the administration of anti-DJ-1 antibody decreased the expression of IL-23 in infiltrating immune cells on day 1 after stroke onset, leading to the suppression of TNFα and IL-1β expression in ischemic brain tissue on day 3 and to neuroprotection against ischemic brain injury on day 7 after stroke onset.

We discovered the previously unknown function of extracellular DJ-1 as a DAMP. DJ-1 could thus be a therapeutic target to prevent excessive inflammation and neuronal injury after ischemic stroke. In addition, DJ-1 expression has been reported to increase in cancer cells; this increase is important for cancer pathology (Kawate *et al*, 2017; Kim *et al*, 2005). Increased expression of DJ-1 within the brain cells has also been observed in several neurodegenerative diseases, such as Parkinson’s disease, amyotrophic lateral sclerosis, Creutzfeldt-Jakob disease, Huntington’s disease, and Alzheimer’s disease (Choi *et al*, 2006; Lev *et al*, 2009; Saito *et al*, 2014; Sajjad *et al*, 2014; Tahir *et al*, 2018). Extracellular DJ-1 may affect the pathologies of cancer and neurodegenerative diseases and could also be a therapeutic target against inflammatory diseases and tissue injuries.

## Materials and methods

### Mice

*DJ*-*1*-deficient mice (*Park7*^−/−^) were purchased from The Jackson Laboratory (Goldberg *et al*, 2005). TLR2 and TLR4 double-deficient mice were kindly provided by Professor Shizuo Akira of Osaka University, Japan. All mice were on a C57BL/6 background. All experiments were approved by the ethics committee and the animal research committee of Tokyo Metropolitan Institute of Medical Science (approval No. 17001).

### Middle cerebral artery occlusion model

A mouse model of transient MCAO was induced by means of an intraluminal suture as described previously (Shichita *et al*, 2012). A >60% reduction of cerebral blood flow was confirmed by laser Doppler flowmetry, and head temperature was kept at 36 °C with a heat lamp. Sixty min after MCAO, the brain was re-perfused by withdrawing the intraluminal suture. Male mice aged (8-10 weeks) were used for all MCAO experiments. 200 μg of control IgG or anti-DJ-1 antibody were intravenously administered from retro-orbital venous sinus immediately after the induction of brain ischemia. To measure infarct volume, each phosphate buffered saline (PBS)-perfused brain was fixed with 4% paraformaldehyde/PBS and embedded in paraffin. A 5 μm section was deparaffinized and stained with anti-MAP2 antibody (Sigma-Aldrich, 1:1000 dilution, Cat# M4403) (the details of this procedure are described elsewhere (Shichita *et al*, 2009)). The infarct area was defined as the MAP2-negatively stained area. Infarct volume was calculated as (contralateral hemisphere – (ischemic hemisphere – infarct area)). Neurological deficits in mice were examined and scored using a four-point scale as previously described (Bederson *et al*, 1986). To investigate the expression of inflammatory cytokines, infiltrating immune cells collected by Percoll-gradient centrifugation (GE Healthcare) or ischemic brain tissues were lysed with RNAiso Plus (Takara). Purified total RNA was transcribed to cDNA using High-Capacity cDNA Reverse Transcription Kit (Applied Biosystems) with random primers. Quantitative PCR was performed using SsoFast EvaGreen Supermix (Bio-Rad) on a CFX96 Real-Time System device (Bio-Rad) with different sets of quantitative PCR primers (**Table S3**). To deplete macrophages from mice, 100 μL of clodronate liposomes (FormuMax Scientific Inc.) was intraperitoneally administered 3 h before MCAO.

### Preparation of recombinant proteins

Complementary DNA encoding candidate proteins identified by liquid chromatography - mass spectrometry (LC-MS) analysis (described elsewhere (Shichita *et al*, 2012)) were cloned from a murine brain cDNA library. cDNA constructs were inserted into the pGEX6P-3 plasmid (GE Healthcare) and expressed as GST-fusion proteins in High efficiency BL21(DE3) Competent Cell (GMbiolab). GST-fusion proteins were purified using glutathione-Sepharose 4B columns (GE Healthcare). Protein-bound glutathione beads were thoroughly washed with cold PBS six times, and thereafter were eluted with 20 mM reduced glutathione (recombinant GST or GST-fusion peptides) or incubated with PreScission Protease (GE Healthcare) overnight at 4 °C to remove the GST tag. These recombinant proteins were incubated with Affi-Prep Polymyxin Media (Bio-Rad) for 4 h at 4 °C to remove endotoxins and endotoxin-bound proteins. To generate modified DJ-1 proteins, which lack the ability to form its oxidized state, recombinant DJ-1 protein was treated with 1 mM dithiothreitol and incubated with 55 mM iodoacetamide (Wako Pure Chemical Industries) in a dark chamber at room temperature for 1 h. We examined the purity of these recombinant proteins by SDS-PAGE with Coomassie brilliant blue (CBB) staining. Recombinant GST protein was used as a negative control for cytokine induction in BMMs.

### Generation of rabbit polyclonal antibody

Rabbits were immunized with recombinant DJ-1 protein. N-hydroxysuccinimide (NHS)-Sepharose beads (GE Healthcare) were crosslinked with recombinant proteins according to the manufacturer’s instructions and were used for the purification of antibodies against DJ-1 protein. IgG antibodies were purified from rabbit serum using protein A sepharose beads (GE Healthcare), and these IgG antibodies were administered to brain ischemia model mice as a control antibody. The specificity of the anti-DJ-1 antibody thus generated was confirmed by Western blotting analysis of lysates from ischemic brain tissue. Sample brain tissues were weighed and brain lysate samples were prepared by homogenizing brain tissue with lysis buffer. 20 μg of brain lysate was examined by SDS-PAGE and proteins were blotted on PVDF membrane (Immobilon, Merck). Membrane was blocked with 2% skim milk and subsequently incubated with anti-DJ-1 antibody overnight. Membrane was washed with PBST and incubated with secondary antibody for 1 h. Designated proteins were detected by Chemi-Lumi One (Nacalai Tesque). Chemiluminesence signal was detected using a Luminescent Image Analyzer LAS-3000 (Fuji).

### Primary bone marrow-derived macrophage culture

The femur and tibia were removed from each mouse and both ends of each bone were cut off. Bone marrow was forced out into 10% RPMI-1640 and the suspension was filtered with a 40 μm filter. Red blood cells were lysed with 17 mM Tris and 48 mM NH_4_Cl and centrifuged for 5 min at room temperature. To generate BMMs, the obtained bone marrow cells were cultured in RPMI-1640 containing 10% FBS and 10 ng mL^−1^ mouse macrophage colony-stimulating factor (M-CSF, Peprotech) in a humidified 5% CO_2_ atmosphere at 37 °C for six days. The cultured BMMs were collected with dissociation buffer (10 mM EDTA/PBS) and were used for subsequent experiments. For experiments with BMMs activated by DAMPs, BMMs were cultured with 1 μM of recombinant GST (control) or 0.1–1 μM DJ-1 protein. The mRNA expression levels were examined by quantitative PCR. TNFα and IL-23 expression in the culture medium was examined using an ELISA (Invitrogen for TNFα Biolegend for IL-23).

### NF-κB luciferase assay

HEK293 cells expressing murine TLR2 or human TLR4 and CD14/MD2 were purchased from InvivoGen. Cells were cultured in the presence of blasticidin (Nacalai Tesque) and hygromycin B (Cayman Chemical) according to the manufacturer’s instructions. Both HEK293 cell lines were transfected with NF-κB luciferase reporter vector and β-galactosidase gene. For the experiments using transiently murine TLR-expressing HEK293 cells, murine TLR2, TLR3, TLR7, TLR9, and Unc93B1 cDNAs were cloned into the pMX-EF1α-IRES2-Venus expression vector. Unc93B1, NF-κB luciferase reporter vector and β-galactosidase gene were transiently co-transfected with each TLRs vector to HEK293 cells. To examine the response to DJ-1 overexpression, the DJ-1 cDNA vector that was cloned into pcDNA3.1 was co-transfected with Unc93B1, NF-κB luciferase reporter vector, β-galactosidase gene, and each of the TLRs vectors. Recombinant DJ-1 protein (1 μM) was added to the transfected HEK293 cell lines and luciferase activities were measured using a Luciferase Assay System (Promega). 10 μg mL^−1^ of peptidoglycan (Sigma-Aldrich), 1 μg mL^−1^ of lipopolysaccharide (Sigma-Aldrich), 5 μg mL^−1^ of Imiquimod (TCI), 10 μM of CpG-DNA, and 10 mg mL^−1^ of poly(I:C) (InvivoGen) were used as positive controls for these experiments. A plasmid containing the β-galactosidase gene was used to normalize for transfection efficiency.

### Immunohistochemistry

For terminal deoxynucleotidyl transferase-mediated dUTP-biotin in situ nick end labeling (TUNEL) staining, paraffin-embedded sections were deparaffinized and incubated with proteinase K (20 μg ml^−1^) and 0.3% hydrogen peroxide. Terminal deoxynucleotidyl transferase and dUTP-biotin were added to each section and the sections were incubated at 37 °C for 1 h. Horseradish peroxidase (HRP) or iFluor 546 (AAT Bioquest, 1:300 dilution, Cat# 16958) was used for the detection of dUTP-biotin. TUNEL-positive neuronal cells were counted in the peri-infarct region, which was considered to extend 3 mm lateral from the midline as previously described (Takada *et al*, 2005). The counts of positive neuronal cells within three different areas (each 0.1 mm square) were expressed as an averaged value. For the detection of extracellular DAMPs, deparaffinized sections were incubated with proteinase K (20 μg ml^−1^) and were blocked with Blocking One Histo (Nacalai Tesque). Sections were incubated with primary antibody overnight at 4 °C and washed with PBS three times. Anti-NeuN antibody (Millipore, 1:500 dilution, Cat# MAB377), anti-F4/80 antibody (Bio-Rad, 1:300 dilution, Cat# MCA497RT), anti-Olig2 antibody (Millipore, 1:300 dilution, Cat# AB9610), and anti-GFAP antibody (Sigma-Aldrich, 1:1000 dilution, Cat# G9269) were used for the detection of neuron, macrophage, oligodendrocyte, and astrocyte, respectively. Anti-pan-cadherin antibody (Abcam, 1:30 dilution, Cat# ab6529) was used for the detection of cellular membrane. Alexa488-conjugated secondary antibody (Thermo Fisher Scientific, 1:300 dilution, Cat# A11070 for rabbit IgG) or Alexa546-conjugated secondary antibody (Thermo Fisher Scientific, 1:300 dilution, Cat# A11081 for anti-rat IgG, Cat# A11018 for anti-mouse IgG, Cat# A11071 for anti-rabbit IgG) was used for the detection of primary antibody. To quantify extracellular DJ-1-positive areas (outside the cellular membrane detected by pan-cadherin staining) or intracellular DJ-1-positive areas (within NeuN-positive cells), each DJ-1-positive area in 50 um square area was measured using Fiji software (NIH). Images of the sections were observed and captured under a fluorescence microscope (BZ-X710, Keyence) or a confocal laser microscopy (LSM710, Carl Zeiss).

### Preparation of infiltrating immune cells from brain

Mice were transcardially perfused with ice-cold PBS to exclude circulating blood cells. The forebrain was homogenized in RPMI-1640 and treated with type IV collagenase (1 mg mL^−1^, Sigma) and DNase I (50 μg mL^−1^, Sigma) at 37 °C for 45 min. Infiltrating immune cells were collected from the interphase of 37/70% Percoll (GE Healthcare) and used for further experiments. For FACS analysis, infiltrating inflammatory cells from ischemic brain tissue were prepared using Percoll and surface staining was performed for 15 minutes with the corresponding mixture of fluorescently-labeled antibodies. Infiltrating inflammatory cells were detected using the following surface markers: microglia included CD45-intermediate and CD11b-intermediate; macrophages included CD45-high, CD11b-high, F4/80^+^, and Gr-1^−^; neutrophils included CD45-high, CD11b-high, and Gr-1^+^. FACS analysis was performed on a FACS Aria III instrument (BD Biosciences) and analyzed using FlowJo software (Tree Star).

### Neuronal cell death assay

Culture supernatant of activated BMMs treated with 1.0 μM of recombinant GST or DJ-1 protein for 6 h was added to the culture of the neuronal cell line, Neuro2A. Neuronal cell death was examined using SYTOXgreen (Thermo Fisher Scientific) at 24 h after adding recombinant proteins or culture supernatants.

### Data Availability

Each coding sequence used for cDNA cloning into expression vector can be obtained from NCBI (gene accession number: *Park7*(DJ-1), NM_020569; *Tlr2*, NM_011905; *Tlr3*, NM_126166; *Tlr7*, NM_001290756; *Tlr9*, NM_031178; *Unc-93b1*, NM_001161428). Crystal structure of DJ-1 protein can be found in Protein data bank (PDB ID: 1P5F).

### Statistical analysis

Data are expressed as means ± s.e.m. One-way ANOVA followed by post hoc multiple-comparison tests (Dunnett’s correction) was used to analyze differences among three or more groups of mice or samples. Between two groups of mice or samples, an unpaired Student’s *t*-test was performed to determine statistical significance. Wilcoxon rank sum test was performed to determine statistical significance of neurological deficits. *P* < 0.05 was considered a significant difference.

## Acknowledgements

We thank Dr. K. Kuroda for mass spectrometry analysis of brain homogenates and Dr. R. Sato and Dr. T. Shibata for giving us technical advices for generating endosomal-TLR-expressing HEK293 cells. We also thank Dr. J. Horiuchi for the assistance to write this manuscript and Dr. H. Shitara and Dr. R. Ishii for supporting animal breeding. This work was supported by the Leading Graduates Schools Program, “Global Leader Program for Social Design and Management” by the Ministry of Education, Culture, Sports, Science and Technology (K.N.), PRIME from AMED under grant number JP20gm5910023 (T.S.), a Grant-in-Aid for Scientific Research on Innovative Areas (Dynamic regulation of brain function by Scrap & Build system) (19H04765) and (Inflammation Cellular Sociology) (20H04957) from the Ministry of Education, Culture, Sports, Science and Technology of Japan (MEXT) (T.S.), JSPS KAKENHI Grants-in-Aid for Young Scientists (17H05096) (T.S.), (18K14831) (S.S.) and (17K15204) (J.T.), a Toray Science and Technology Grant (T.S.), the Takeda Science Foundation (S.S.), and by grants from the Mitsubishi Foundation (T.S.), the SENSHIN Medical Research Foundation (T.S.), the MSD Life Science Foundation (T.S.), the Senri Life Science Foundation (T.S.), and the Ono Medical Research Foundation (T.S.).

## Author Contributions

K.N. designed and performed the experiments, analyzed the data, and wrote the manuscript; S.S. and J.T. provided technical advice; A.N., K.O. participated in data analysis and discussion, Y.Y. and K.K. provided experimental supports and performed the experiments; H.M. provided technical advice about experimental design; T.S. initiated and directed the entire study, designed experiments, and wrote the manuscript.

